# Continuous land cover change detection in a critically endangered shrubland ecosystem using neural networks

**DOI:** 10.1101/2021.10.26.465837

**Authors:** Glenn R. Moncrieff

## Abstract

Existing efforts to continuously land cover change using satellite image time-series have mostly focused on forested ecosystems in the tropics and northern hemisphere. The notable difference in reflectance that occurs following deforestation allows land cover change to be detected with relative accuracy. Less progress has been made in detecting change in low productivity, disturbance-prone vegetation such as grasslands and shrublands, where natural dynamics can be difficult to distinguish from habitat loss. Renosterveld is a hyperdiverse, critically endangered shrubland ecosystem in South Africa with less than 5-10% of its original extent remaining in small, highly fragmented patches. I demonstrate that direct classification of satellite image time series using neural networks can accurately detect the transformation of Renosterveld within a few days of its occurrence, and that trained models are suitable for operational continuous monitoring. A dataset of precisely dated vegetation change events between 2016 and 2021 was obtained from daily, high resolution Planet labs satellite data. This dataset was then used to train and evaluate 1D convolutional neural networks and Transformers to continuously detect land cover change events in multivariate time-series of vegetation activity of Sentinel 2 satellite data as it becomes available. The best model correctly identified 89% of land cover change events at the pixel-level, achieving a f-score of 0.93, a 79% improvement over the f-score of 0.52 achieved using a method designed for forested ecosystems based on trend analysis. Models have been deployed to operational use and are producing updated detections of habitat loss every 10 days. There is great potential for supervised approaches to continuous monitoring of habitat loss in ecosystems with complex natural dynamics. A key limiting step is the development of accurately dated datasets of land cover change events with which to train machine learning classifiers.

## Introduction

Land cover change is the single largest cause underlying the global biodiversity crisis (Brondizio et al., 2019). It is estimated that at least 21% of all terrestrial plant extinctions are directly related to agriculture alone (Le Roux et al., 2019). A core component of efforts to halt biodiversity loss and climate change are remote sensing-based monitoring systems built upon satellite-derived earth observation data (Turner, 2014; Turner et al., 2003; Yang et al., 2013). These systems allow up-to-date reporting on trends in habitat loss, and in some instances, near-real time alerts that may aid interventions and enforcement. Typically, temperate and tropical forests are the focal ecosystems for applications, examples of which include Global Forest Watch and Terra-i.

While forests are indeed globally important stores of carbon and home to myriad species, far less attention has been paid to developing remote sensing-based monitoring for use in open ecosystems where trees may be present but not dominant, such as savannas, grasslands, shrublands and deserts. Open ecosystems cover a large proportion of the globe, make up 40% of the global total ecosystem organic carbon and harbour a substantial proportion of the world’s biological diversity (Bond, 2019; Poulter et al., 2014)). Failure to develop capacity for change detection in open ecosystems would exclude the majority of the earth’s terrestrial surface from monitoring. Detecting abnormal change in forested ecosystems is relatively simple in comparison to open ecosystems. The spectral signature of a closed-canopy forest stands in stark contrast to the sparsely vegetated or bare ground that remains after deforestation. In contrast, the natural vegetation state in open ecosystems can vary dramatically due to natural disturbances, long-term trends or cyclical functions, such as those relating to fire, seasonality and interannual weather variations (Slingsby et al., 2020; Wilson et al., 2015). This variability found within healthy ecosystems can obscure spectral change that occurs when intact natural vegetation is converted to nonnatural land use.

The Cape Floristic region (CFR) in South Africa is home to some of the world’s most floristically diverse open ecosystems (Manning and Goldblatt, 2012; Myers et al., 2000). Within the CFR the most threatened vegetation type is the grassy shrubland known as Renosterveld (Skowno et al., 2019, 2021). Renosterveld is restricted to relativity nutrientrich soils in the south and west of the Western Cape Province. It is noteworthy for its exceptionally high diversity of bulbs and - due to widespread conversion of intact Renosterveld to agriculture - perhaps the highest concentration of plants threatened with extinction in the world (Brummitt et al., 2015; Humphreys et al., 2019; Raimondo, 2011; Skowno et al., 2021). Only 5-10% its original extent remains, with much of this contained in small isolated fragments (Von Hase et al., 2003). Despite legislation prohibiting the destruction of Renosterveld habitat, loss is ongoing (Moncrieff, 2021). It is estimated that at current rates Renosterveld will become extinct as an ecosystem within the 21st century (Skowno et al., 2021). The primary cause of this decline is the expansion of intensive agriculture (primarily for the cultivation of grains and oil-seed such as wheat, barely and canola), and overgrazing by livestock.

The current approach to conservation in the region is clearly inadequate. A remote sensing-based monitoring system, such as is available for many forest ecosystems worldwide, may help to reduce rates of Renosterveld loss by aiding in the identification of land owners responsible for historical loss - thereby deterring future habitat destruction. If information is provided with low latency between the onset of ploughing and its detection, there may be potential to intervene while habitat destruction is underway. Continuous change detection approaches attempt to assimilate and analyze new remote sensing data as they become available, and have the potential to provide near-real time alerts on land cover change. A typical pattern used for continuous change detection involves analyzing trends in historical vegetation activity and building a model that can accurately forecast the expected reflectance of intact forests. Predictions are then compared recently acquired observations (Verbesselt et al., 2012; Ye et al., 2021a; Zhu and Woodcock, 2014; Zhu et al., 2020). When observed vegetation activity falls outside the range expected for natural vegetation, this is taken to be indicative of forest loss. Trend analysis methods used to model expected vegetation activity typically assume a relatively predictable progression of vegetation activity in intact ecosystems that can be well described by linear models with harmonic terms to capture seasonality (Zhu and Woodcock, 2014). The resulting land cover change detections, while impressive for stable ecosystems such as forest, are prone to commission errors in dynamic ecosystems where natural variability can easily be mistaken for land cover change and omission errors when disturbance is subtle. These problems are particularly prevalent in non-forest ecosystems with natural dynamics such as fire, aridity and herbivory (Slingsby et al., 2020).

When ground-truth data that distinguish areas where habitat loss has occurred are available, land cover change can be directly classified from time-series of vegetation activity using supervised learning. This removes the need to specify a model for expected vegetation activity (Hansen et al., 2013; Kennedy et al., 2009). Hansen et al. (2016) trained decisiontree classifiers to detect forest disturbance in the tropics, comparing recent imagery to a stable reference period. Their system is widely used for forest monitoring in the tropics and has been incredibly impactful. In Renosterveld, Moncrieff (2021) used a training dataset of validated land cover change events to produce a map of habitat loss between 2016 and 2020 after manually verifying potential change identified through model predictions. These approaches are however limited to producing updates on land cover change at discrete intervals (often annually), as the training data typically do not specify the exact date on which change occurred, but rather that change occurred at some unknown point between two fixed points in time (e.g. January and December). In this example a new full year of input data from January to December is required before a new prediction can be made. While this provides useful information for reporting on habitat loss, this information is usually only obtained months or years after the reported change has occurred (Hansen et al., 2013; Kennedy et al., 2010).

If a precise date for a land cover change event is available, supervised methods can be trained to classify change using input data covering a period of time shorter than year (Figure 1). Furthermore, if these labelled events occur at a variety of points in time throughout the year (and potentially across multiple years) models that can recognise the signal of land cover change regardless of the time of year and are robust to intra and inter-annual natural variability can be trained (Pacheco-Pascagaza et al., 2022). These models can then be applied continuously as new data become available and produce near-realtime land cover change alerts. They would also be less prone to errors than change detection methods requiring the ecosystem to follow a predictable pattern (e.g Verbesselt et al. (2012); Zhu and Woodcock (2014); Zhu et al. (2020)), as the models could theoretically learn to distinguish the spectral patterns indicative of land cover change from natural disturbances at any point in time if shown sufficient examples.

**Fig. 1.**
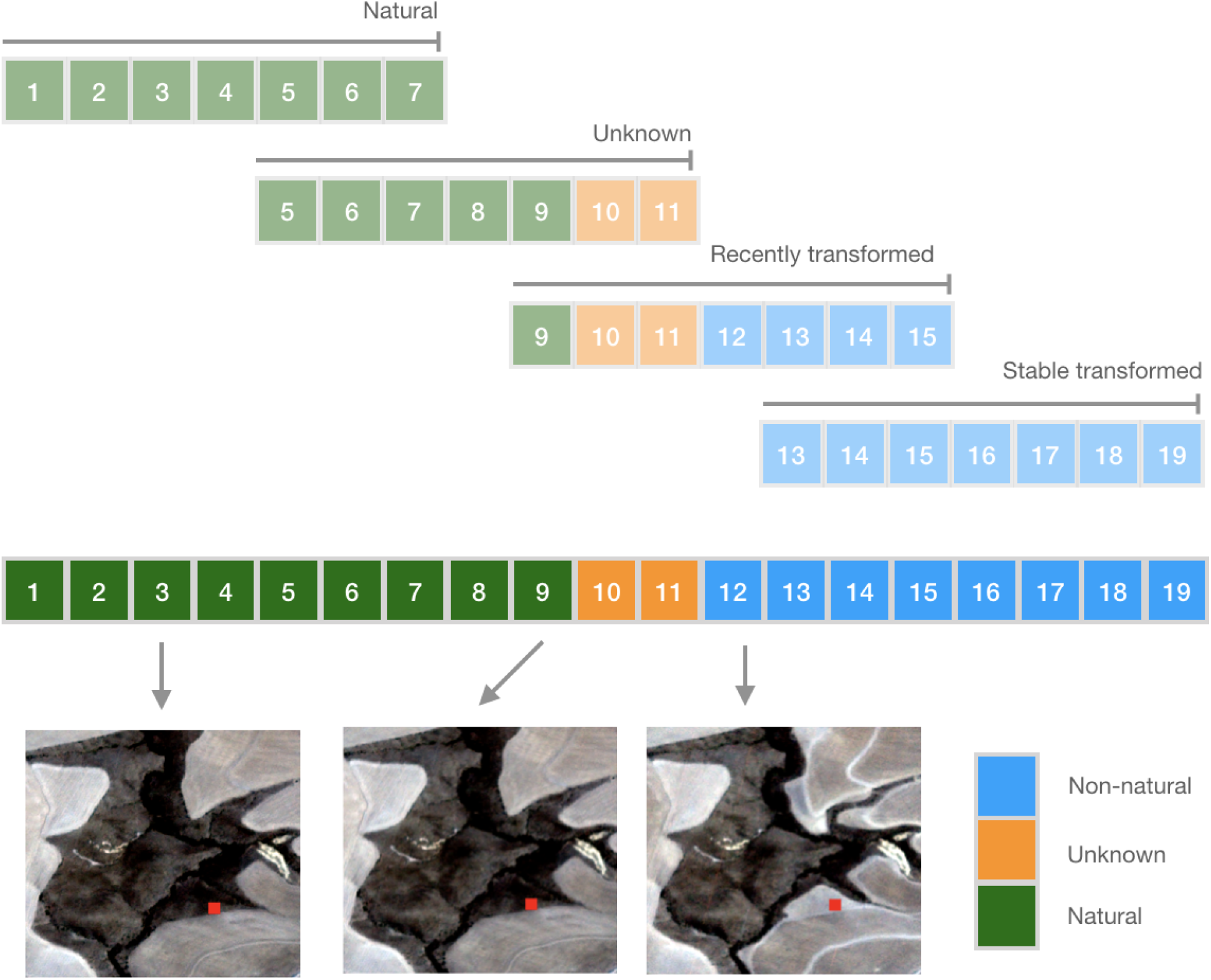
The approach used for labelling time series to enable continuous classification of change. Each step in the sequence of 19 observations represents an observation obtained for a single pixel at a regular interval (e.g. 10 days). The land cover label for the pixel (identified by the red point in the images) can either be natural, unknown or non-natural. This label is obtained through inspection of time series’ of Planet Labs PlanetScope satellite imagery. For the example pixel indicated here, an image obtained at time step 3 indicates that the land cover is natural on this date, the image at time step 9 indicates the last date on which natural land cover is observed. The first time at which non-natural land cover is observed is step 12. Hence, it is not known whether the land cover was natural or non-natural at timestep 10 and 11. Using a window length of 7 observations and forward step of 4, 4 labelled sequences can be extracted from the full time series. Each sequence is assigned the label of the most recent observation. Provided enough labelled events occurring at different points in time across multiple years, models that can recognise the signal of land cover change regardless of the time of year and are robust to intra and inter-annual natural variability can be trained. These models can then be applied continuously as new data become available to produce near-realtime land cover change alerts.

Machine learning approaches such as random forests and support vector machines have been widely used to directly classify land cover change using multi-temporal remotely sensed input data (Habib et al., 2009; Hansen et al., 2013; Mas, 1999; Wessels et al., 2016). These methods do not however explicitly take into account the temporal structure within the data. Neural networks specifically designed for classification of sequential data such as convolutional neural networks (CNN) or Transformers have been shown to improve performance on time series classification tasks relative to other machine learning methods (Ismail Fawaz et al., 2019). They have recently been being applied to land cover classification with great success (Kussul et al., 2017; Pelletier et al., 2019; Rußwurm and Körner, 2019) and direct classification of land cover change (Sefrin et al., 2020).

Here I describe the design and performance of a system for continuous land cover change detection in a critically endangered shrubland ecosystem, the Renosterveld of South Africa. Using a dataset of precisely dated land cover change events covering multiple years, two different deep learning architectures are trained and evaluated against tree-based and trend analysis methods commonly used for land cover change detection and monitoring in forested ecosystems. Model building and design choices are motivated by an intention to produce a system that can be deployed for operationally use by conservation management and law enforcement agencies. This demands a system that is robust to the natural variability within this ecosystem, can assimilate and make predictions on new data as it becomes available, and is able to detect habitat loss with minimal delay.

## Methods

### A. Study region

The study region is limited to the extent of the Renosterveld bioregion in the Overberg district municipality in the Western Cape Province of South Africa (Mucina and Rutherford, 2006). The choice of region was determined by the critically endangered status of Renosterveld vegetation, the availability of accurate land cover data for the Overberg municipality, and partner conservation organizations in the region. The Overberg municipality covers an area of of 12,241 km^2^, with Renosterveld covering 4711 km^2^ with this region naturally. Detailed mapping of remaining natural vegetation in the Overberg based on 2003 data identified 656 km^2^ of the original extent of Renosterveld vegetation in the Overberg remaining.

### B. Land cover change events

Using random forests for change detection on Sentinel 2 imagery and subsequent detailed verification with very high resolution imagery, Moncrieff (2021) identified all conversion of Renosterveld to nonnatural land cover between January 2016 and January 2020. Land cover change was identified within 268 parcels (a spatially contiguous land cover change event) totalling 478.6 ha. Using daily Planet Labs PlanetScope data, dates were assigned to each event indicating the closest possible timing for the occurrence of the land cover change event. It is not possible to assign every event to a single day as often multiple days or even weeks elapse between the beginning and end of a land cover change event. Furthermore, clouds and haze can reduce the frequency at which clear observations are available. Thus for each land cover change event two dates were assigned - the latest date on which the presence of natural vegetation could be confirmed, and the earliest date on which the absence of natural vegetation could be confirmed (Figure 1). These two dates occur within one week for 21% of all events, and within two months for 87%. Here a sample of these data are used where land cover changes dates were assigned with the highest confidence, leaving 185 events and 219 ha of land cover change.

This dataset contains only events in which complete conversion of intact natural vegetation occurred and transformation into non-natural land cover could be dated within a short period of time. It thus excludes gradual degradation occurring over multiple months or years. Fire is a natural process in Renosterveld and is not considered a land cover change event unless it precedes further modification and subsequent conversion to croplands. An additional 4 events covering 7 ha were identified within the same region between Jan 2020 and Jan 2021. These events were used for testing model performance outside of the temporal range of training data, providing a more accurate assessment of operational performance. To augment data with additional observations of stable natural vegetation, parcels of vegetation that remained natural over the entire study period were added. This added an additional 2590 ha dataset covering the period 2016-2019 and 24 ha to the testing dataset covering 2020. These no-change parcels were selected from the (Von Hase et al., 2003) map of remaining Renosterveld, buffered to ensure that they did not occur within 100m of non-natural vegetation and manually verified.

### C. Sentinel 2 time series

Input time series’ for classification were created using Sentinel 2 L1C data (S2). For training and validation all available S2 data up until 2019-12-31 were extracted and preprocessed, whereas testing data covered the period 2020-01-01 to 2020-12-31. S2 images are pre-screened and discarded if total scene cloud cover exceeds 50 percent. Pixel-level cloud masking is performed using the s2cloudless machine learning-based cloud masks. Pixels with a cloud probability greater than 40 percent are masked. Cloud shadows are also masked and are calculated from cloud masks using cloud and solar geometry combined with dark pixel detection. All S2 bands are retained and the following additional vegetation indices calculated: Normalized Difference Vegetation index (NDVI), Enhanced Vegetation Index (EVI), Normalized Difference Water index (NDWI), Normalized Burn index (NBR) and Normalized Difference Red-edge index (NDRE). Data are exported at 10m resolution with 20m and 60m bands resampled using bilinear interpolation.

S2 data area available every 10 days in this region until mid-2017 when data becomes available at least every 5 days subsequent to the launch of Sentinel 2B. In parts of the study region data is available at frequencies higher than 5 days due to swath overlap. To create regular time series from the irregularly spaced S2 image stacks pixel-level time series are created with a single observation every 10 days. Where multiple images are available within a 10 day period, data from the image with the highest pixel-level NDVI value are used. Where no data is available, the reflectance and vegetation indices from the previous valid image is used unaltered. In this application interpolation between observations by averaging is undesirable as it may obfuscate sudden changes in reflectance diagnostic of land cover change. Classification is performed on time series of reflectance collected over 180 days, equalling 18 observations with a 10 day interval. The choice of 180 days was arrived upon through experimentation with both longer and short time series’. While it is theoretically possible to detect Renosterveld loss with as few as 2 sequential observations if the change occurs rapidly, few of the labelled events have such precision in determining the actual data on which habitats loss occurred. Longer time series contain more information on the spectral properties of vegetation before and after change, but no additional performance was gained when experimenting with time series longer than 180 days.

### D. Data labelling

The S2 time series for each pixel within a parcel is assigned the label ‘natural’ if the last date of confirmed natural vegetation falls after the end date of the time series under consideration (Figure 1), or if no change was ever reported within that parcel. If both the last date of confirmed natural vegetation and the first date of confirmed land cover change occur within the 180 day time series’ the land cover change event must have occurred within the time series under consideration and the ‘recently transformed’ label is assigned. If the last date of confirmed natural vegetation occurs within the time series and the first date of confirmed land cover change falls after the end of the time series it is unclear whether land cover change has indeed taken place and the data point is discarded. Similarly, if the first date of confirmed land cover change occurs before the start of the time series, land cover change has already taken place. Here we do not include a stable non-natural land cover class, thus these data are discarded.

The resulting labelled dataset is highly imbalanced, with the vast majority of sampled pixels in the ‘natural’ class. There is also very high spatial auto-correlation among neighbouring pixels within the same parcel. In addition, successive time series for the same pixel obtained 10 days apart will share 94 percent of observations. To ameliorate class imbalance data and spatial autocorrelation only 5 percent of pixels within parcels where no change occurs are sampled and 33 percent of pixels from parcels where change does occur are used. To reduce temporal oversampling within a pixel, time series windows of 180 days are shifted forward by 60 days for every sample taken. This represents a trade off between reducing temporal oversampling and discarding labelled data. Subsequent to this data thinning, pixels are stratified by parcel and assigned to training and validation sets using a 70/30 split - that is 70 percent of parcels are allocated to the validation set and 30 percent to the validation set. This ensures that data from within the same parcel cannot be split between the training and validation sets and mitigates the impact of spatial autocorrelation on performance evaluation (Wadoux et al., 2021). Test data is derived from land cover change events in the year following the period over which training and validation data was derived. This is intended to evaluate model performance in an operational setting and hence does not undergo temporal or spatial thinning, with a new prediction made every 10 days.

### E. Models

Two neural network architectures that have shown excellent performance in classification of land cover from satellite image times series are trained and evaluated along with random forests, a tree based method commonly used for land cover classification. TempCNN is a 1D convolutional neural network for land cover classification from satellite image times series (Pelletier et al., 2019). Convolutional kernels that extract relevant features in local temporal regions across multivariate time series of individual pixels are learned. These 1D CNN layers are followed by non-linear activations and potentially also batch normalization and stacked, before class probabilities are produced by a dense and softmax layer. TempCNN performs well on a range of land cover classifications tasks, and has been adopted by multiple packages for classification of satellite image time series (Rußwurm et al., 2020a; Simoes et al., 2021).

Transformer models were originally proposed as sequenceto-sequence encoder-decoder models based on the self-attention operation and applied to language translation (Vaswani et al., 2017). Attention mechanisms operate by assigning an importance-score (attention) to weight each element in a sequence (Bahdanau et al., 2016). In self-attention, the concept of attention is used to encode sequences to a hidden representation. In the traditional application of transformers to language tasks, self-attention encoders are followed by self-attention decoders. For time series classification only the encoder network is needed. The encoder network learns which elements in a time-series are relevant for the classification task at hand and extracts salient features. In this work, a Transformer model that has been evaluated in Rußwurm and Körner (2019) is used, where individual pixels’ time series’ are encoded using stacked self-attention layers. This architecture outperforms many other neural networks and tree-based methods for crop type mapping using satellite image time series (Rußwurm et al., 2020a).

### F. Model fitting

Hyperparameter optimization for random forests, TempCNN and the Transformer network is performed using a grid search over 24, 36 and 32 parameter combinations respectively. The best fitting model of each class is then evaluated against test data. For the Transformer and TempCNN, the Adam optimizer is used is minimize focal loss. TempCNN and Transfomers as implemented in Tensorflow 2 and the eo-flow python package. scikit-learn is used for Random Forests. Google Earth Engine is used for preprocessing of input data (Gorelick et al., 2017).

To aid model interpretation, the most important temporal regions identified by the TempCNN model for detecting land cover change are calculated using Grad-CAM++ (Chattopadhay et al., 2018). Originally developed for use with CNNs and image classification, Grad-CAM++ can easily be used when CNNs are applied to time series classification. Grad-CAM++ calculates a saliency score (importance) for each time step in a time series toward the final prediction made by the model. This is calculated using a weighted average of all positive activation gradients across channels in the final convolutional layer.

### G. Trend analysis comparison

The accurately dated land cover change data used here to train supervised methods is seldom available. Trend analysis approaches to land cover change detection are more generally applicable and widely used (Bullock et al., 2020; Tang et al., 2019). A method based on trend analysis is used to evaluate the improvement in accuracy achieved when using direct classification on these data. The approach followed is very similar to the approached outlined by Zhu and Woodcock (2014), termed Continuous Change Detection and Classification (CCDC). CCDC is widely used for continuous change detection, and implemented in platforms such as Google Earth Engine. More recent methods based on similar principles that would improve on this baseline are available (Ye et al., 2021a,b; Zhou et al., 2019), but they are not commonly used on S2 data.

The CCDC approach operates by fitting a simple linear model with a single harmonic term to capture seasonal variation using robust regression through iteratively reweighted least squares with the Talwar cost function. The model is fitted to historical NDVI data for every pixel in regions known to still contain Renosterveld. The expected NDVI over the test period (2020-01-01 - 2020-12-31) is then calculated for each day of the year at each pixel. This is compared to the observed NDVI and anomalies calculated and expressed in terms of the RMSE of the fitted regression. If 3 successive observations differ from the prediction by 3 times the RMSE, land cover change is assumed to have occurred. The RMSE thresholds used to trigger detection can be changed in order to account for ecosystems with greater inherent variability (Zhu and Woodcock, 2014). Different thresholds were experimented with, with 3 times the RMSE providing the best performance here. More sophisticated implementations of this workflow make use of multiple bands, or use parameters learned from labelled land cover change events (Ye et al., 2021b).

## Results

Full time series’ of selected pixels extracted for the entire period covered by the training and validation data are shown in Figure 2. These data show the highly dynamic and variable nature of vegetation in this ecosystem. In natural vegetation NDVI peaks in winter, corresponding to the wet season, but the onset of greening is highly variable. Summer minimum NDVI is very low, though both minimum and maximums are highly variable across the region. Nonetheless a distinctive signal attributable to land cover change is visible, and models are able to use this signal to reliably detect land cover change. The best TempCNN model used a dropout rate of 0.8, L2 kernel regularization with a regularization factor of 1e-8, He normal kernel initialization, 6 stacked convolutional layers each with 32 filters and 256 neurons in the final fully connected layer. The best performing Transformer used a dropout rate of 0.5, 4 layers in the encoder with 4 attention heads, a model depth/query size of 128, no layer normalization and 256 neurons in the encoder feed-forward layer. The best performing random forest model used 100 estimators, and required a minimum of 4 samples to split an internal node.

**Fig. 2.**
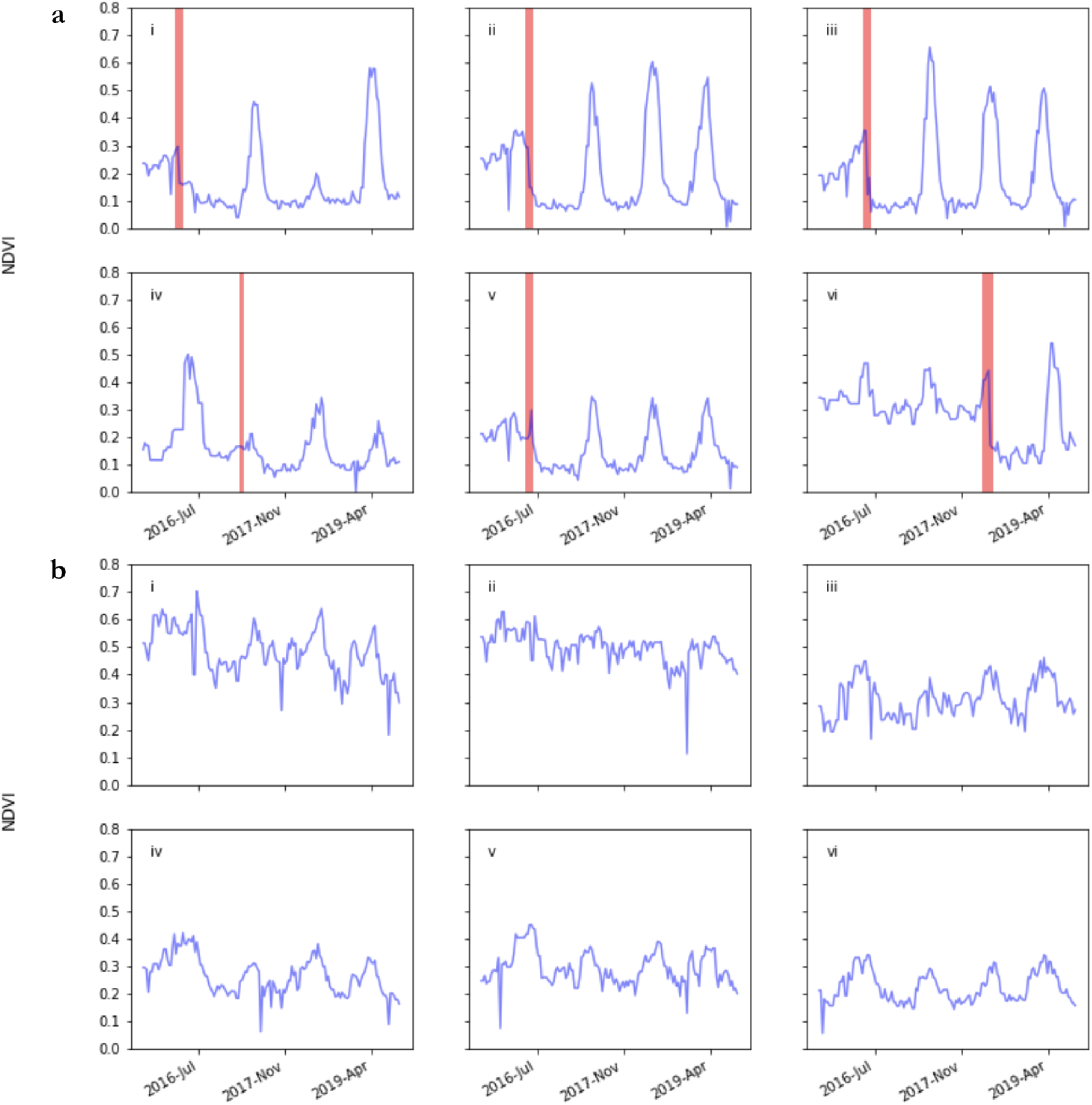
Observed NDVI time series for the full period covered by training and validation data for randomly selected pixels from across the study region. Vegetation in all time series’ begins as natural Renosterveld. In panel a, a land cover change event occurs in the period highlighted in red, and thereafter the land cover is non-natural. In b, vegetation remains natural for the entire time series

The TempCNN achieved the highest performance, with a recall of 0.89 and precision of 0.96 when identifying recently transformed Renosterveld and overall f-score of 0.93 (Table 1). The Transformer was only slightly worse than the TempCNN with a recall of 0.88 and precision of 0.97 on the recently transformed class and overall f-score of 0.92, while the Random Forest was significantly worse than both TempCNN and the Transformer, with recall and precision of 0.79 and 0.98 on recent transformation, and overall f-score of 0.88. All models using direct classification outperformed the trend analysis method, which was far more likely to produce both false negatives and false positives with recall of 0.49 and precision of 0.54 on the recently transformed class and overall f-score of 0.52.

**Table 1.** Model performance evaluated on test data. Recall-1-short reports model recall when change has occured within the first 2 time-steps (20 days), of the 18 step (180 days) time series. Performance of the top-performing model for each metric is highlighted in bold.

|  | Accuracy | F1 | Recall-1 | Recall-0 | Precision-1 | Precision-0 | Recall-1-short |
| --- | --- | --- | --- | --- | --- | --- | --- |
| Random Forest | 0.98 | 0.88 | 0.79 | **0.99** | **0.98** | 0.98 | 0.50 |
| TempCNN | **0.99** | **0.93** | **0.89** | **0.99** | 0.96 | **0.99** | **0.64** |
| Transformer | **0.99** | 0.92 | 0.88 | **0.99** | 0.97 | **0.99** | 0.60 |
| CCDC | 0.90 | 0.52 | 0.49 | 0.95 | 0.54 | 0.94 | - |

Performance for random forests, TempCNN and the Transformer declines when their ability to detect recent transformation within 20 days of change is measured. Again the TempCNN performed best, with recall of 0.64, followed by the Transformer with 0.60 and the random forest with 0.50. Trend analysis classification is not possible within 20 days of change, as 3 anomalous data points are required before change is flagged. This equates to 30 days with the 10 day interval used here.

Figure 3 provides a visual depiction of the time series features characteristic of Renosterveld loss events, determined using Grad-CAM++ on the trained TempCNN. For a single Renosterveld loss event in a pixel viewed from multiple points in time after the event, the same temporal features are focused upon. Across multiple different land cover change events, the network has learnt to consistently focus on time periods in which a drop in vegetation greenness has occurred. Other more nuanced changes in spectral reflectance may also be contributing to the importance of these temporal regions, as the TempCNN is using all input bands and indices to classify change, though only NDVI is depicted here.

**Fig. 3.**
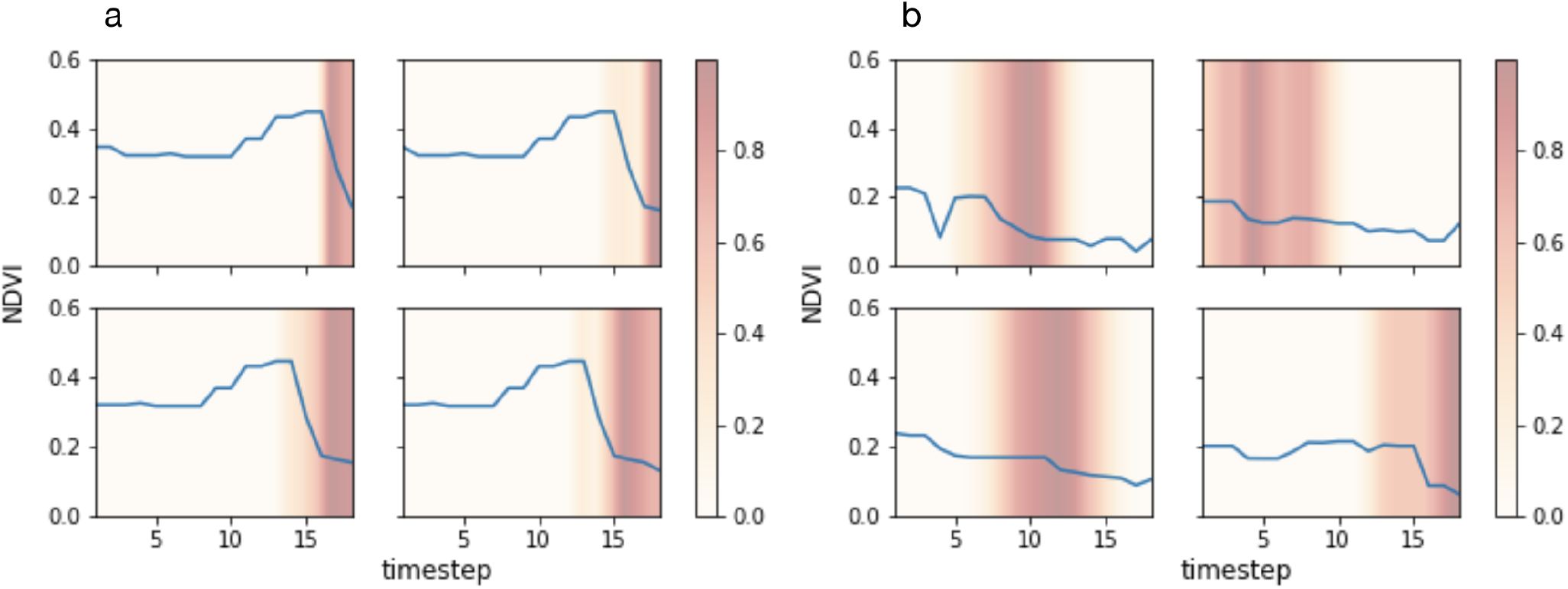
Saliency of temporal regions in time series for pixels in which land cover change is predicted to have occurred, determined using Grad-CAM++. Higher saliency scores indicates that the region is more influential in determining that land cover change has occurred. Blue lines show NDVI, though saliency is calculated using all input features. In a) a single pixel is shown at four points in time, with each subsequent panel shifted forward 1 step (10 days). In b) time series’ from four spatially and temporal independent pixels are shown.

A prototype application performing up-to-date inference over the entire study region every 10 days using TempCNN is available at https://glennwithtwons.users.earthengine.app/view/globalrenosterveld-watch and an example sequence of predictions over 2 months in an area through the course of a land cover change event is depicted in Figure 4. Inference over the entire remaining distribution of Renosterveld in the Overberg region for a single date reveals multiple as-yet unreported areas where Renosterveld has been transformed to cropland within the 6 month time series of satellite data under consideration. Because models are trained to only make predictions in regions where land cover is intact Renosterveld at the beginning of the time series under consideration, predictions are not reliable when made on other vegetation types, or where Renosterveld has already been converted to agriculture. This is not an issue when up-to-date land cover maps are available. Multiple observed cases of incorrect predictions of Renosterveld loss where land cover has in reality remained as agriculture throughout the time series under consideration highlight the need for updating of existing regional land cover maps.

**Fig. 4.**
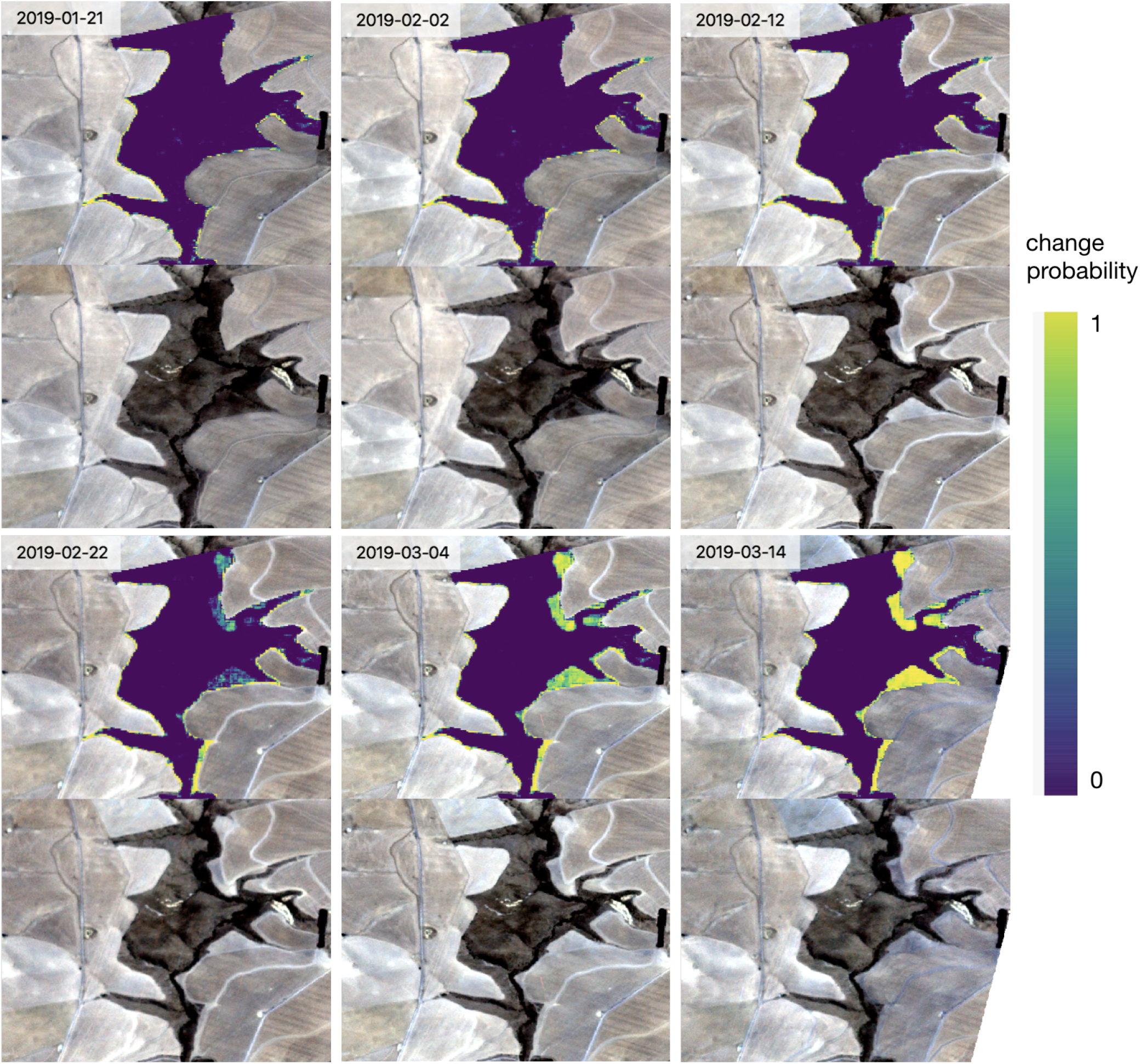
Example sequence of predictions through the course of a land cover change event in Renosterveld shrublands of the Overberg region, South Africa. Upper panels show the predicted probability of land cover change, where predictions are only made in pixels known to have intact natural vegetation at the start of the monitoring period. The lower panels show the most recent Planet labs PlanetScope imagery collected prior to, or on the date for which predictions are made. The detection of Renosterveld loss along the eastern edge of the monitored area is apparent. This detection occurs within 10-20 days of the actual land cover change event.

## Discussion

Shrublands are globally important repositories of biodiversity and play an important role in regulating planetary biochemical cycles (Bond, 2019; Poulter et al., 2014). In many regions shrublands are being lost at a rate that exceeds other ecosystems such as forests (Parr et al., 2014). Despite this, few studies have attempted to develop algorithms designed specifically for operational monitoring of shrubland loss, ostensibly because many of these ecosystem display complex natural dynamics and have low overall productivity. These characteristics make the methods typically applied in operational forest monitoring perform poorly. This study demonstrates that reliable detection of land cover change in shrublands in possible, and that detection is possible within days of land cover change events, thus facilitating the use of these algorithms for operational monitoring and enforcement. The key component of this is the continuous use of direct classification, enabled by the availability of hand-labelled change events with the date of each event precisely determined.

Direct classification is rarely used for continuous detection of land cover change. The labels needed to train models require detailed examination of very-high resolution satellite or aerial imagery and dense time series. Very-high resolution data may not be available for the required region, time period or frequency, and the costs associated with these will be prohibitive for many potential use cases. Even when all the necessary data is available, the data labeller must have intimate knowledge of the ecosystem in question in order to determine whether an observed event is indeed land cover change. For example, Renosterveld is a naturally fire-prone ecosystem and burnt vegetation is not necessarily indicative of land cover change. These limitations should not, however, preclude the development of labelled datasets. The availability and cost of very-high resolution and high temporal frequency imagery is improving dramatically, and new programmes are broadening access to these data (e.g. the Norway International Climate and Forest Initiative - Planet partnership). Advances in deep learning performance on land cover classification when labelled data are limited will further reduce data requirements (Rußwurm et al., 2020b; Sun et al., 2021). The investment of time required to acquire labels is indeed significant, but for critically endangered ecosystems that are predicted to become extinct within decades based on current trajectories, surely this investment is justifiable. Particularly if there is reason to believe that an operational land cover change monitoring system will slow or reverse these trends.

The volume of labels required to achieve a performant model in the example presented here is surprisingly small - just 219 ha of dated land cover change events. This is sufficient to produce large improvements in performance in deep learningbased classifiers relative to random forests which typically require less data to train. While it is highly unlikely that a supervised approach will be applicable in any region beyond the domain of the training data (thus requiring further painstaking data collection and labelling), meta-learning and transfer learning approaches ought to reduce data requirements for application to new disturbance types and new geographies (Vanschoren, 2019). Given multiple labelled datasets of dated land cover change from different geographies or ecosystems, models can be trained such that they are initialized optimally and learn rapidly when confronted with a new regime. Particularly promising is model-agnostic metalearning, which does not necessitate custom designed models for this task, but rather can make use of widely adopted models such as TempCNN or Transformers (Finn et al., 2017; Rußwurm et al., 2020b).

The models developed here are not only limited in their spatial domain of applicability, but also the types of land cover change events which they are capable of detecting. Only a single land cover change can be identified the sudden and destructive conversion of natural Renosterveld to non-natural land cover. Other types of change can occur in this landscape, such as gradual degradation through overgrazing or spread of invasive alien plants. It is possible that the same approach could be used to diagnose these types of changes. This would however require a significant increase in the amount of training data and longer input time series, as these types of change occur over periods longer than the 6 month windows input into classifiers. The choice to focus on a single type of rapidly occurring change allowed relatively high performance to be achieved with limited training data and is focused on the actions considered most severe by law enforcement in the region.

A major improvement achieved through the approach presented here is the ability to detect change within a few days of the actual event. Rapid identification is critical in many intended applications of near-realtime continuous land cover change monitoring - such as law enforcement and management intervention. Other approaches typically require multiple consecutive observations confirming the change has occurred before a positive identification is made (Ye et al., 2021a; Zhu and Woodcock, 2014). Generally any land cover change will be complete by the time these detections are reported. Here change is identified with relative skill with only 2 observations subsequent to the actual change event. In certain applications this may be sufficient to lead to an onthe-ground intervention that could preventing further damage. Accuracy improves as further observations are acquired, and hence it is possible to delay the identification of change depending on the cost/benefit of any particular application. The improvement in performance obtained when using deep learning-based time-series classifiers vs. random forests is likely a result of their ability to retain temporal information and extract features based thereon. Previous work comparing the performance of Transformers and TempCNN for land cover classification found Transformers to generally perform better than TempCNN, particularly when minimal prepossessing and no atmospheric correction is performed (Rußwurm and Körner, 2019; Rußwurm et al., 2020a). Here, extensive feature engineering has been conducted and cloud masking performed to remove noisy data, potentially resulting in the convergence in performance between TempCNN and the Transformer. A range of machine learning methods that are not based on neural networks have been developed for the classification of time-series data, and there may be potential performance gains when using these methods, particularly when training data is scarce (Dempster et al., 2020; Maus et al., 2016). Gains in performance cannot, however, come at the costs of increases in model inference time to the extent that prediction cannot be made at the required latency or within available budgets. This is particularly relevant because monitoring will often be required over very large regions. Using the TempCNN model fitted here prediction over the entire remaining range of Renosterveld in the Overberg region can be achieved within 120 CPU hours. The full prediction pipeline for preprocessing, prediction and postprocessing is achieved at minimal cost using tools such as Google Earth Engine and Google Cloud Dataflow.

## Conclusions

The potential of continuous change detection using deep learning on satellite image time series, particularly in nonforest ecosystems, remains largely unexploited. This study demonstrates that when labelled data are available, accurate and precise detection of land cover change is possible at low latency. Obtaining the labelled data does indeed require significant effort, and it is likely that independent datasets and models for will be required for different ecosystems. This should not discourage the further development and operationalization of this approach. Both deep learning and the explosion of remote sensing data have been lauded as having great potential to address the biodiversity and climate crises. For these technologies to truly have impact they must find application beyond the forested ecosystems of the world, and embrace the complexity of open ecosystems.

## ACKNOWLEDGEMENTS

I am very grateful for the support and encouragement given by Odette Curtis and her passion for conservation in this region. Marcel Gietzmann-Sanders helped produce the prototype operational monitoring application. Multiple individuals from the Western Cape Department of Environmental Affairs and Development Planning are acknowledged for their input, in particular, those from the Environmental Law Enforcement directorate. This research was supported by the National Research Foundation of South Africa through (Grant No. 118593) as part of the RReTool: Rapid and repeatable tools for monitoring and mitigating global change impacts on natural resources project, and the Group on Earth Observations-Google Earth Engine Programme. The funders had no role in study design, data collection and analysis, decision to publish, or preparation of the manuscript.

## DATA AVAILABILITY

Code and data used in this analysis are available at https://github.com/GMoncrieff/renosterveld-monitor. Due to ongoing criminal investigations location data has been removed to protect the identity of landowners where necessary.

